# Where the magnet sits sets the sign of the scaling law: membrane-anchored versus intracellular magnetic actuation of cells

**DOI:** 10.64898/2026.11.10.765704

**Authors:** Yutian Xia, Lifeng Yin, Angelina Lin, Joanna Tang, Zhaoqing Cong

## Abstract

Magnetic nanoparticles apply mechanical force to cells in two incompatible geometries: anchored to the plasma membrane, where a rotating field twists a surface-bound cap, and internalized into lysosomes, where an assembled rod stirs the organelle lumen. They are treated as interchangeable, and the choice is made on synthetic convenience. They obey opposite design rules. Treated analytically at matched single-cell magnetic moment, the step-out frequency of an intracellular rod falls with assembly size while that of a membrane cap rises linearly; the scaling exponents have opposite sign and the curves cross at 184 particles and 4.04 Hz, inside the window in which published experiments operate. The cap tension resultant reduces to an identity in which cell radius cancels, leaving magnetic texture as the decisive design variable: a field-oriented cap reaches 2.69 mN m^−1^, the same particles with random easy axes reach 0.0033 mN m^−1^, and a superparamagnetic cap transmits no torque at all. The model reproduces four published measurements with no fitted parameters. One disagrees: our intralysosomal wall shear stress is 33 to 81 times below the published value while reproducing every structural feature of it. The model yields two zero-predictions that one experiment would falsify.

## 1 Introduction

A magnetic nanoparticle on or in a cell is a mechanical actuator with an unusual property: the operator switches it on and off at a distance, without changing anything else about the cell’s environment. Two constructions dominate. In the first, particles are anchored to the plasma membrane, often as a Janus cap on one hemisphere, and a rotating field twists the cap against the bilayer and the underlying cortex [1]. In the second, particles are internalized, accumulate in lysosomes, assemble under the field into short rods, and stir the organelle lumen [2,3,4]. Each descends from a distinct lineage. The membrane route grew out of work that binds particles to surface receptors or channels and then pulls or twists them, from selective activation of mechanosensitive channels [5] to receptor clustering by nanomagnetic actuation [6] and death-receptor aggregation by magnetic switching [7], surveyed early as remote control of cellular behavior [8], and extended to lithographically defined spin-vortex microdiscs interfaced with the cell surface [9]. The intracellular route grew out of particles bound to or enclosed within an internal membrane and then spun or oscillated until that membrane leaks [10,11,12]. Both are described as ways of applying force to a cell, and the choice between them is usually presented as a matter of synthesis.

We argue that the choice is the single largest determinant of what the experiment does, because it sets which mechanosensitive structure the delivered force can load. A cap on the outer leaflet loads the plasma membrane and the cortex, where tension-gated channels and cortical tension sensors sit [13,14]. A rod inside a lysosome loads the limiting membrane of an acidic organelle, where the readouts are membrane permeabilization, galectin recruitment and cathepsin leakage [15,16] [17,18]. Macrophages read cyclical force through PIEZO1 [19], and the transcriptional machinery that sustained force engages, the YAP/TAZ route and the lamin-linked nuclear route, is characterized mainly in other cell types [20,21]. These are different structures with different thresholds, and nothing guarantees that loading one reproduces the cell-state change that loading the other produces. The published record already contains two experiments pointing in opposite directions: a driven intralysosomal assembly produces durable M1-like macrophage repolarization [2], while a very-low-field extracellular swimmer suppresses the PIEZO1–AP-1–CCL2 axis and moves macrophages towards M2 [22].

Our hypothesis is that placement, not dose, is the primary design variable. At matched single-cell magnetic moment the two routes deliver stimuli that differ in magnitude, in frequency dependence and in the identity of the structure loaded, so that the direction of the biological response is an output of the placement decision rather than a property of magnetic stimulation as such.

Treating the response direction as an output is what makes the argument testable. If it is a premise, the two contradictory experiments are an embarrassment to be explained away. If it is an output, they are two data points the model must cover, and the model is falsified by the cells it fails to cover.

Reported forces attributable to magnetic actuation span six decades, and each is stated in whichever unit its source preferred, so the record cannot be read as a single scale (figure 1*a*). It is also not equally informative about the two routes, and this asymmetry shapes what can be modeled. We assembled every physical constant entering either arm, with its source, its locator in that source, and a classification of whether it was measured, inferred, assumed, or computed here (electronic supplementary material, table S1). Of 142 entries, 63 are measured values traceable to a figure, table or body-text statement in a primary source, 38 are inferred under an assumption the source does not state, 30 are computed here, two are assumed outright, and **two have never been measured at all**: the rupture tension of the lysosomal limiting membrane, and the in-plane tension screening length of the plasma membrane. Both sit on the critical path (figure 1*b*).

**Figure 1.**
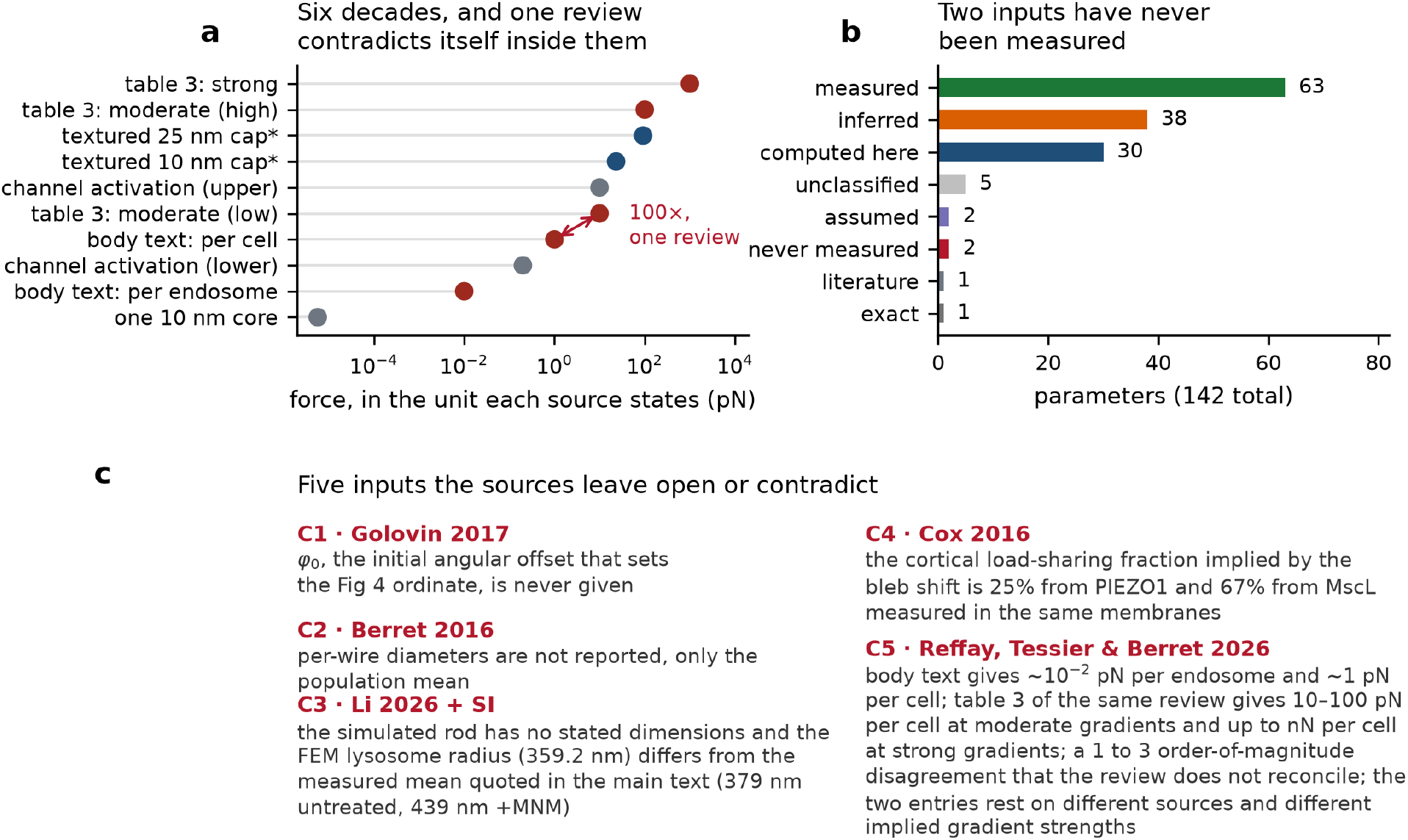
The published record is asymmetric and contradicts itself in five places. **(a)** Forces attributable to magnetic actuation, each plotted against the unit in which its source states it, spanning six decades. Blue, membrane-anchored constructs; dark red, internalized constructs; gray, reference bands. The arrow marks the 100-fold disagreement between the body text and table 3 of a single review (C5). **(b)** Classification of the 142 physical inputs that enter either arm (Supplementary Table S1). Two inputs have never been measured: the lysosomal rupture tension and the plasma-membrane tension screening length. **(c)** The five registered source contradictions, recorded rather than resolved.

**Figure 2.**
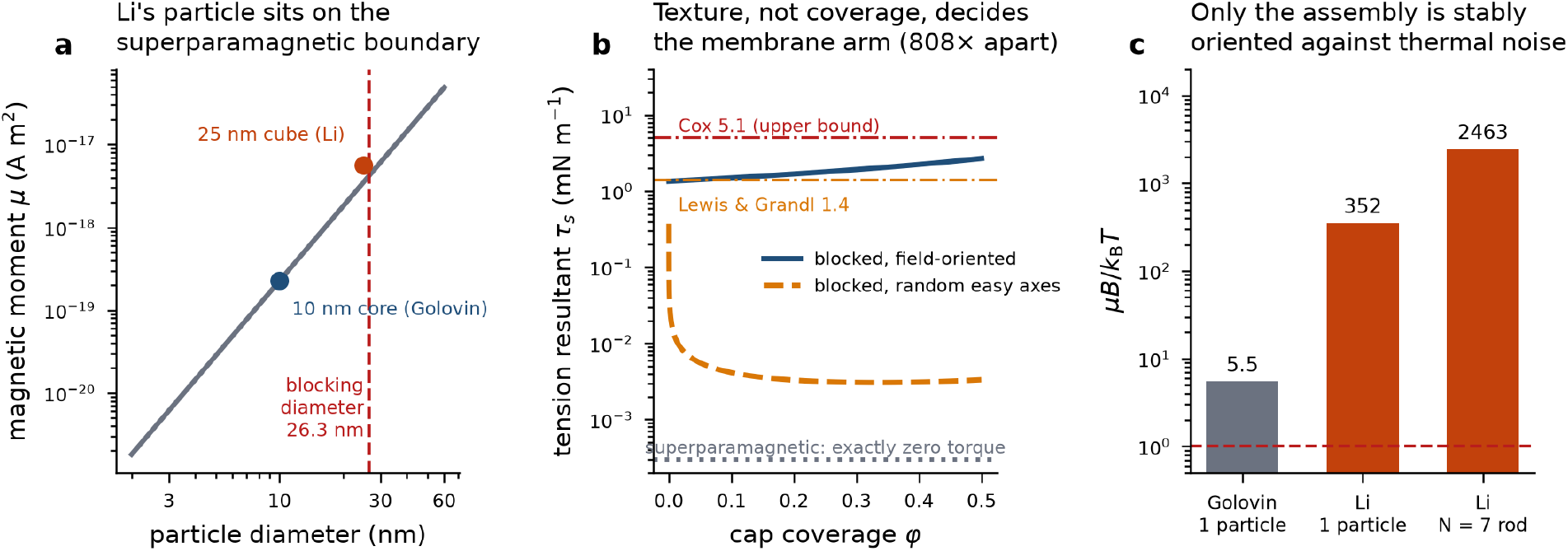
The two routes do not start from the same kind of particle. **(a)** Single-particle moment versus diameter from material constants. The superparamagnetic blocking diameter for magnetite at 300 K (*τ*_m = 100 s) is 26.3 nm; the 25 nm cube sits on the boundary. **(b)** Cap tension resultant versus coverage at 0.26 T for 25 nm cubes under three magnetic textures, against the two published PIEZO1 thresholds. The superparamagnetic case is identically zero and is drawn at the axis floor. **(c)** Magnetostatic energy relative to thermal energy at the field each source uses.

The asymmetry is not uniform. The intracellular arm inherits a characterized particle, a measured lumenal viscosity and a measured organelle radius. The membrane arm inherits a particle and then nothing: no measured cap coverage, no measured anchor density, no measured cortical load-sharing fraction for the construct in question, and no measured screening length. The two arms are not equally constrained by data, and a comparison that does not say so is comparing a measurement with an extrapolation.

Five source inputs are internally contradictory or absent (figure 1*c*; electronic supplementary material, table S3). The initial angular offset that sets the ordinate of Golovin and co-workers’ figure 4 is never given [23,24] (C1). Berret reports only a population median wire diameter although the methods state each wire was measured individually [25] (C2). Li and co-workers’ simulated assembly has no stated dimensions, and their finite-element lysosome radii, 359.2 and 453.0 nm, differ from the measured means quoted in their main text, 379 and 439 nm [2] (C3). Cox and co-workers’ cortical load-sharing fraction is 25% from PIEZO1 and 67% from MscL in the same membranes [13] (C4). Reffay and co-workers’ body text gives about 1 pN per cell while table 3 of the same review gives 10–100 pN at moderate gradients and up to nN at strong ones [26] (C5). Each is recorded with the handling we adopt rather than resolved.

One claim of novelty, and only one. Intralysosomal magneto-mechanical actuation is not new [2,3,4], nor is magnetically induced lysosomal membrane permeabilization [17], nor the galectin readout of magnetically induced endosomal damage [18]. A citation-network walk over a two-step neighbourhood of 831 works around seven seeds, extended by twelve targeted searches returning 396 further works, found no prior work that models the two placements side by side at matched magnetic moment, and none that treats response direction as a model output.

## 2 Model

Full derivations, parameter provenance and the frozen validation criteria are in the electronic supplementary material. The essentials follow.

### Magnetic moments

come from material constants with no adjustment. A 10 nm magnetite core at the saturation magnetization and density used by Golovin and co-workers [23] carries *µ* = 2.262 ×10^−19^ A m^2^; the 25 nm zinc-doped ferrite cube used by Li and co-workers [2] carries *µ* = 5.606× 10^−18^ A m^2^. The superparamagnetic blocking diameter for magnetite at 300 K with a 100 s measurement time is 26.3 nm [27], so the 25.0 nm cube sits on that boundary (figure 2*a*) and the magnetic texture of an assembly is treated as a free design variable rather than a given. At the field each source uses, magnetostatic energy exceeds thermal energy by a factor 5.5 for a 10 nm core at 0.1 T, 352 for a single 25 nm cube at 0.26 T and 2463 for a seven-cube assembly, so thermal randomization of an assembled rod is never the limiting consideration (figure 2*c*).

### Hydrodynamic drag

Rod rotational friction is computed on two calibers in parallel, because they differ by a factor 1.5 [28] and the paper should not pretend otherwise: the Broersma slender cylinder [29] gives *ξ* = 1.870 ×10^−21^ N m s for seven cubes end to end in lysosomal fluid, the Chwang and Wu prolate spheroid [30], the form used by the source we validate against, gives *ξ* = 1.241× 10^−21^ N m s. Whole-cell drag is the Stokes sphere, *ξ* = 8*πη*R^3^ = 1.060 ×10^−17^ N m s at R = 7.5 *µ*m. **The whole cell is 5670 times harder to turn than the rod it contains**, and that single ratio drives most of what follows. Wall traction inside a spherical cavity is the exact axisymmetric rotlet solution *τ*(*θ*) = 3T sin *θ*/(8*π*R^3^) with the finite-size correction applied as in the source.

### Viscosity calibers are never interchanged

extracellular medium 10^−3^ Pa s; lysosomal contents of a loaded organelle 0.4711 Pa s; cytosol seen by a micron-scale probe 47 (+26/ −14) Pa s [25], the value our level-2 validation fixes, against 1 to 3 Pa s on nanometer probes [31]. They appear in separate rows of every table.

### Rotation dynamics

Two torque branches are treated separately: a permanent moment, torque∝ *µ*B sin *φ*, and an induced moment, torque ∝*µ*_0_Δ*Χ* H^2^ sin 2*φ*. Mean rotation rates come from numerical integration of the lag-angle equation, checked against the closed form Ω = *ω* − *√* (*ω*^2^ − *ω*_c^2^).

### Membrane arm

The cap is a rigid patch bonded over a spherical cap of half-angle *θ*_c; its torque is reacted as an in-plane stress resultant at the rim, *τ*_s = T/(2*π* a_cap^2^) with a_cap = R_cell sin *θ*_c. Tension propagation is computed both cortex-free and screened over a screening length of 0.1–10 *µ*m.

### Dose

D ∝*τ*^2.416 Δt^0.785 [32], anchored on the plasma-membrane condition, computed independently with the published and with our own lysosomal wall-shear value.

## 3 Results

### 3.1 The two placements have step-out exponents of opposite sign

A magnetically driven rotor follows the field only while magnetic torque overcomes viscous drag. Above a critical drive frequency, the step-out frequency f_c, it desynchronizes and its mean rotation rate collapses. Step-out is the natural currency for comparing placements: it is set entirely by the ratio of the magnetic ceiling to hydrodynamic drag, and both are computable (electronic supplementary material, table S4).

Adding particles is the one design variable an experimenter controls, and the two routes answer it in opposite directions (figure 3*a*). For the **intracellular rod**, particles lengthen the rod; moment grows as N but rotational drag grows faster, roughly as L^3^ with a logarithmic correction, so f_c ∝ln N/N^2^, with a local exponent of −**1.75** at N = 100 that tends to −2 as the rod lengthens. More particles make the intracellular actuator worse at high frequency. For the **membrane cap**, particles grow the moment as N while the load, the cell, is unchanged, so f_c ∝N with a local exponent of exactly **+1.000**. More particles make the membrane actuator better.

**Figure 3.**
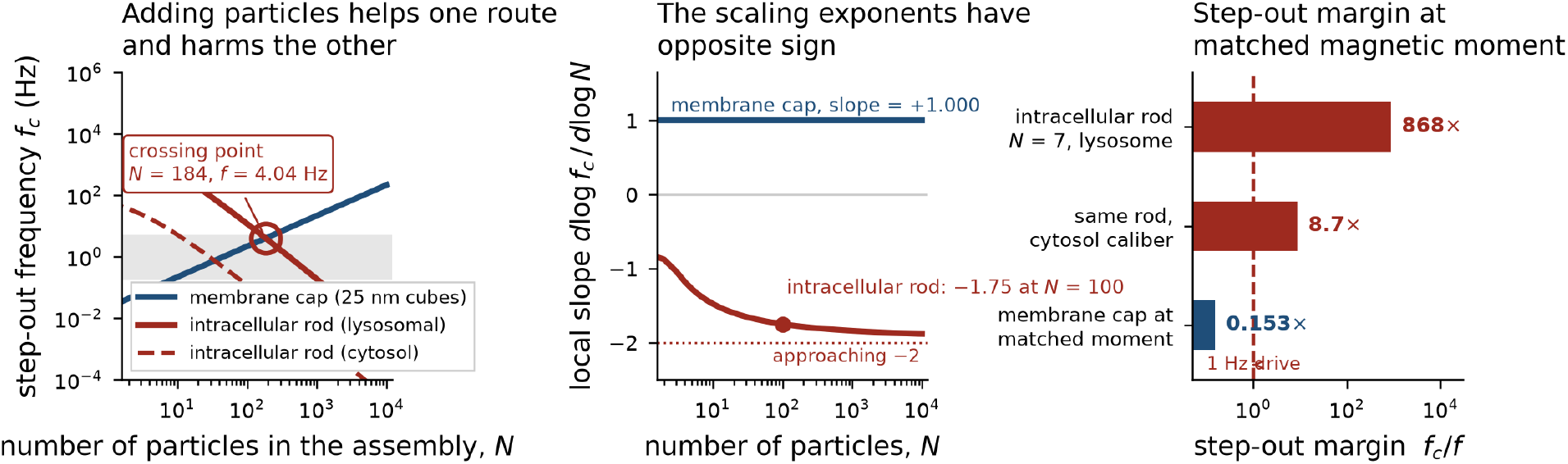
The central result: the two placements have step-out exponents of opposite sign. **(a)** Step-out frequency versus assembly size for the intracellular rod (lumenal and cytosolic viscosity calibers) and the membrane cap. Shading marks the 0.2–5 Hz window in which published intralysosomal experiments operate. The routes cross at N = 184 and 4.04 Hz. **(b)** Local scaling exponent d log f_c/d log N: +1.000 for the cap; for the rod it is−1.75 at N = 100 and tends to−2, as the panel annotates. **(c)** Step-out margin f_c/f at a 1 Hz drive for three matched-moment configurations; the dashed line is the synchronous boundary.

**Figure 4.**
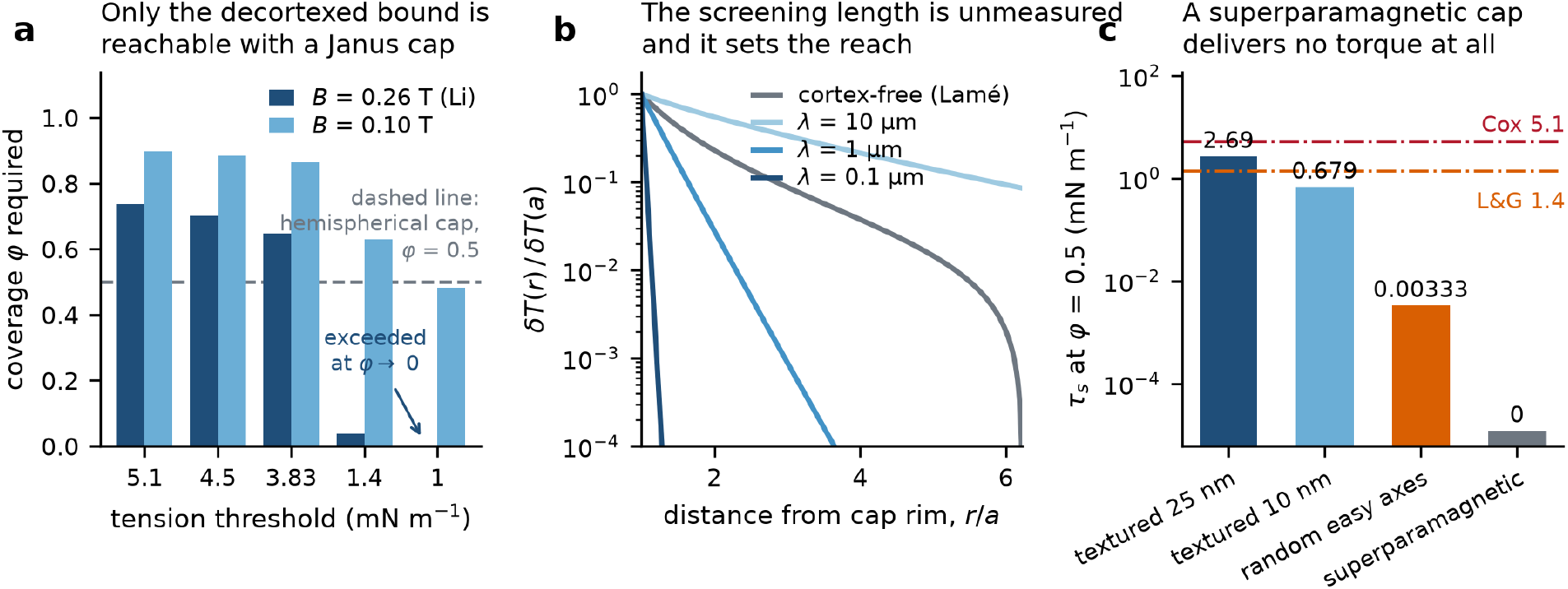
The membrane arm, treated analytically. **(a)** Coverage required to cross each tension threshold, at two field strengths, for 25 nm cubes. **(b)** Tension propagation away from the cap rim for the cortex-free solution and for screening lengths spanning the plausible range; the prediction changes qualitatively across it, and *λ* has never been measured. **(c)** Tension resultant at *φ* = 0.5 u*√*nder three magnetic textures, the textured case shown at both particle sizes; the random-easy-axis penalty is *N* = 808-fold and the superparamagnetic case is exactly zero.

The exponents have opposite sign (figure 3*b*), and the curves cross at **N = 184 particles and f = 4.04 Hz** (figure 3*a*), inside the 0.2–5 Hz window in which the published intralysosomal experiments are run. Below the crossing the intracellular route has the larger frequency margin; above it, the membrane route does.

At matched single-cell moment the asymmetry is stark (figure 3*c*). Li’s as-published configuration, a seven-cube rod in a 359 nm lysosome at 0.26 T driven at 1 Hz, has a step-out margin f_c/f of **868**. The same rod released into the cytosol, where the micron-scale caliber is 47 Pa s rather than the lumenal 0.47 Pa s, has a margin of **8.7**: still synchronous, by two decades less. A membrane cap carrying the same moment on the same cell at the same field has f_c = **0.153 Hz** and has therefore already desynchronized at 1 Hz. Giving a cap a comparable 1 Hz margin requires matching the whole-cell intracellular load of about a hundred rods, which raises f_c to 15.3 Hz.

Two corollaries follow. **The viscosity caliber matters more than the organelle size.** Changing the lysosome radius from 359.2 to 453.0 nm leaves f_c unchanged to the printed digits, because step-out is set by the rod’s own drag and not the cavity; changing the medium from lumenal contents to cytosol changes f_c a hundredfold. A rod that escapes its lysosome is a different mechanical object.

#### Adhesion switches the membrane arm between two regimes

The equations of motion give a useful identity: delivered torque as a fraction of the magnetic ceiling equals drive frequency as a fraction of step-out frequency, T/mB = f/f_c. For the intracellular rod at 1 Hz this fraction is **0.115%**. The rod spends almost none of its available torque, because the lumen barely resists. For a cap on an *adherent* cell the rotor cannot turn at all, the torque is reacted entirely by the anchors, and T/mB = 1. Adhesion therefore switches the membrane route from drag-limited to torque-limited, and the intracellular route has no such switch. This is why the same particles at the same field can mean two different mechanical experiments depending on whether the cells were plated or in suspension, a variable published protocols frequently do not state.

### 3.2 The membrane arm: texture, not coverage, is the design variable

Modeling the cap as described and reacting its torque at the rim, the tension resultant reduces to

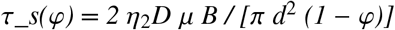

with *φ* the fractional coverage, d the particle diameter and *η*_2_D the areal packing of the monolayer. Three consequences follow.

#### Cell radius cancels

It enters the cap rim radius and the torque and divides out exactly. Cap tension is a material property of the particle layer, not a geometric property of the cell. Cell radius survives only in the step-out frequency, where it enters as R^3^.

#### Tension is almost flat in coverage

Over the Janus range it changes by less than a factor of two, diverging only as *φ→* 1. A bigger cap buys moment and rim circumference in nearly equal proportion, so coverage is not the knob it is assumed to be.

#### Texture is the knob

Figure 2*b* sweeps the tension resultant across coverage for all three textures against both published thresholds, and figure 4*c* is the cut through it at half coverage. At *φ* = 0.5, 25 nm cubes and 0.26 T, a blocked cap with field-oriented easy axes delivers **2.693 mN m**^−1^, which is 0.53 of the Cox upper bound and 1.92 tim*√*es the decortexed lower bound. The same cap with random easy axes delivers **0.0033 mN m**^−1^, a penalty of 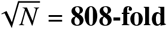, because random moments add incoherently. A superparamagnetic cap delivers **zero** (figure 4*c*). Nothing else in the membrane arm spans five orders of magnitude on a variable the synthesis controls.

Inverting for the threshold (figure 4*a*): crossing the Cox upper bound of 5.1 mN m^−1^ at 0.26 T requires coverage *φ* = 0.736, about 9.6 ×10^5^ particles, which is no longer a Janus cap; the alternative is to raise the field at *φ* = 0.5 to **0.492 T**. The decortexed lower bound of 1.4 mN m^−1^ needs only *φ* = 0.038. A conventional Janus cap at conventional fields therefore sits between the two published tension thresholds, and which side it lands on depends on the cortical load-sharing fraction that C4 leaves contradictory.

The second never-measured quantity enters here. Whether rim stress stays local or propagates depends on the in-plane screening length *λ*, unmeasured for this membrane. Across 0.1–10 *µ*m the predicted stress a few rim radii away changes by more than three orders of magnitude (figure 4*b*), and the qualitative prediction, local blister against whole-cell tension rise, reverses within that range. Direct measurement in other cells shows that tension does not equilibrate instantaneously across the plasma membrane and can stay local over micrometers [33], a reading that remains under discussion [34,35]; none of that work supplies *λ* for a macrophage. *λ* is a bound on the model, not a number we can supply.

### 3.3 Matched-moment comparison

Figure 5*a* ranks thirteen configurations by step-out margin at a common 1 Hz drive, and figure 5*b* recasts the same ranking as the fraction of the magnetic torque ceiling each configuration actually spends. Every intracellular configuration sits in the synchronous regime with margins of 8.7 to 868; every membrane configuration matched to a realistic intracellular load sits at or below the synchronous boundary. Membrane caps climb back above it only when their moment is pushed to the monolayer ceiling, at which point they are no longer Janus caps.

**Figure 5.**
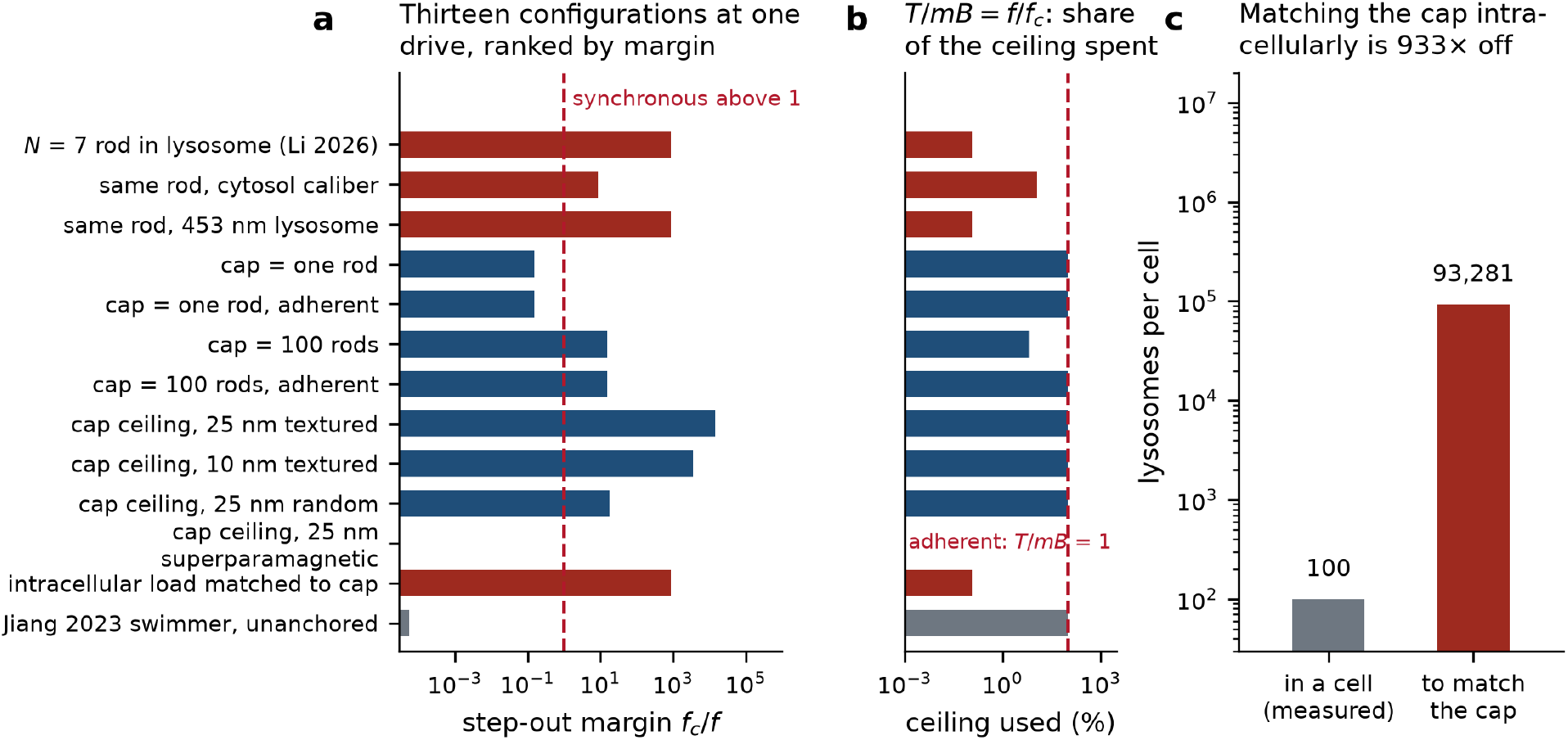
Matched-moment comparison. **(a)** Thirteen configurations ranked by step-out margin at a common 1 Hz drive; dark red, intracellular; blue, membrane-anchored; gray, extracellular unanchored. **(b)** The same configurations as a fraction of the magnetic torque ceiling, using the identity T/mB = f/f_c. Adherent caps sit at 100 % because the rotor cannot turn. **(c)** Iron-loaded lysosomes required to match the moment of a textured 25 nm hemispherical cap, against the literature loading.

The reciprocal statement is sharper. Matching the moment of a textured 25 nm hemispherical cap with internalized particles would require **93,281 iron-loaded lysosomes per cell** against a literature value of roughly 100, a factor of 933 (figure 5*c*). The two routes are not interchangeable at any achievable loading: the membrane route carries far more moment, and for that reason it is the route that saturates the torque ceiling and desynchronizes.

### 3.4 Validation against four published measurements, with no fitted parameters

A model assembled from closed forms earns attention only by reproducing measurements it did not see. We fixed, before running anything, a four-level ladder with pass criteria written down in advance (figure 6; electronic supplementary material, table S2).

**Figure 6.**
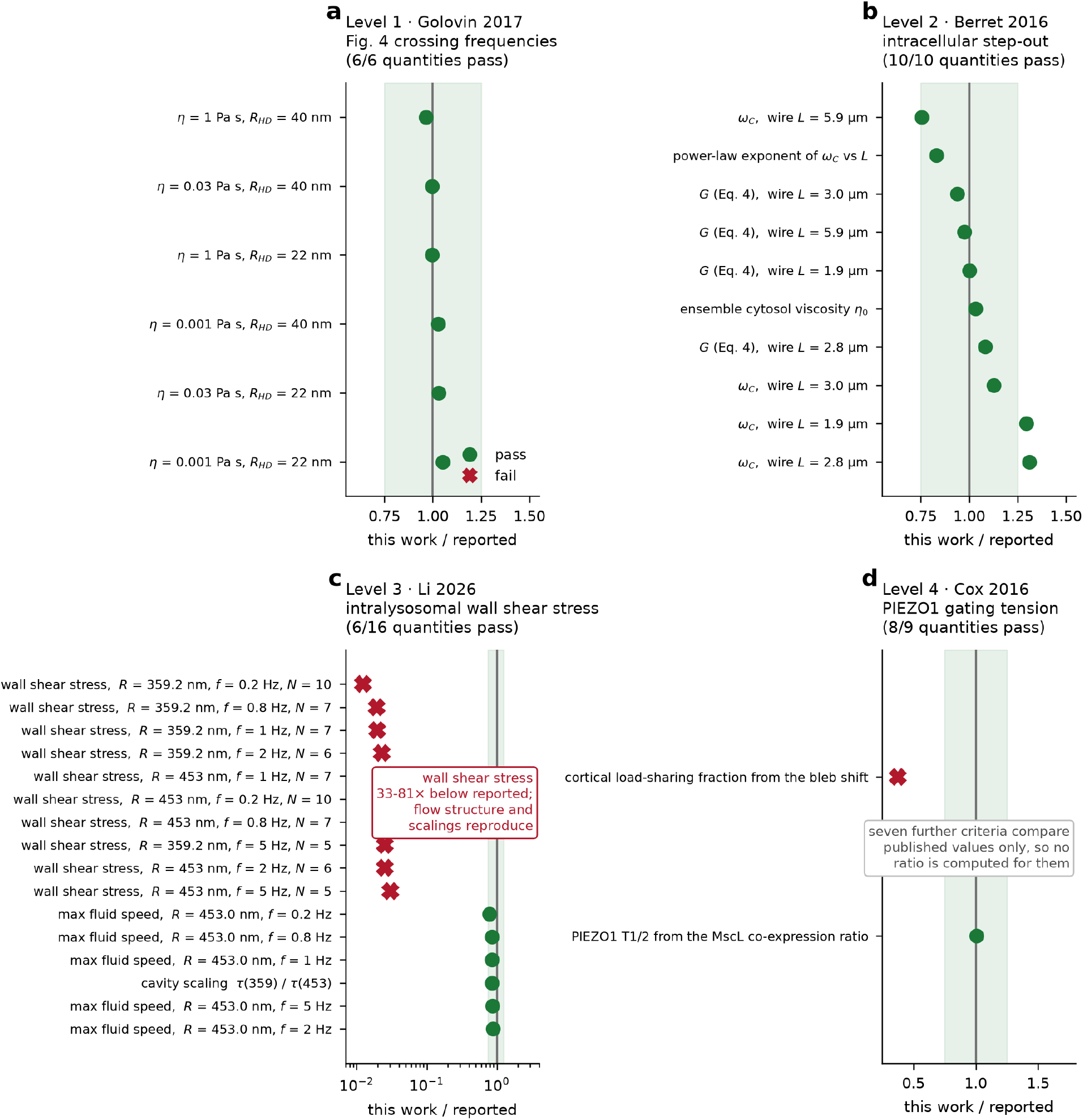
Validation ladder: four published measurements, no fitted parameters. Each panel shows the ratio of our value to the reported value for every quantity at that level for which a ratio can be formed; the shaded band is ±25 %. At level 4 only two quantities yield a ratio, because the remaining seven criteria compare published values with one another; all nine are listed in electronic supplementary material, table S2. Green, criterion passed; red cross, failed. Levels 1, 2 and 4 pass without tuning (the single Level-4 failure is the source’s own internal contradiction, C4). At Level 3 the structural quantities fall inside the band while the wall shear stress is 33-to 81-fold low across ten conditions. Quantities, units and verdicts in Supplementary Table S2.

#### Level 1, Golovin 2017, oscillating field, permanent-moment branch [23]

Six crossing frequencies across three viscosities and two hydrodynamic radii are reproduced within 0.96–1.05 of the reported values. The unreported initial angle (C1) is handled by reproducing the figure caption’s geometry and declaring the inconsistency.

#### Level 2, Berret 2016, rotating field, induced-moment branch [25]

The synchronous-to-asynchronous transition of superparamagnetic wires inside living fibroblasts. Eight pre-registered criteria, all met with no tuning. The forced-exponent fit returns an ensemble cytosolic viscosity of **48.5 Pa s** against the published 47 (+26/ −14) Pa s; the free power-law exponent is −2.49 ±0.58 against the published −3.0±0.5; elastic moduli recovered from the oscillation amplitudes are 16.0, 9.8, 10.3 and 7.8 Pa against the published 16, 9, 11 and 8 Pa; integration of the equation of motion reproduces the closed-form mean rotation rate to 2.6 ×10^−10^; and the asynchronous oscillation frequency tends to 2.000 times the drive, the signature of the induced-moment branch.

Levels 1 and 2 together matter more than either alone, because they exercise *different torque laws*. The blocked lysosomal rod belongs to the permanent-moment branch, superparamagnetic wires and caps to the induced-moment branch. Both are now anchored to published data.

#### Level 3, Li 2026, intralysosomal wall shear stress: structure reproduces, magnitude does not [2]

This level is reported in full because it fails. Eight criteria were pre-registered; six pass, one partially, one fails. Reproduced with no tuning: the monotone rise of wall stress with frequency even as the assembly shrinks from ten to five particles; the greater-than-twofold rise across the frequency range, where we obtain 4.32×; the R^−3^ cavity scaling; strict linearity in viscosity; the rigid-body tip velocity at 0.79–0.88 of the printed maxima; a Reynolds number of 2.6 ×10^−10^; and agreement between two independent torque calculations to within 1.28-to 1.39-fold, inside the factor-of-two criterion. **Not reproduced: the absolute magnitude, 33 to 81 times below the reported values across ten conditions.** At 1 Hz, R = 359.2 nm, 471.1 cP and N = 7 we obtain 0.0201 Pa against the reported 1.03 Pa.

The discrepancy is not a shape detail. Inverting the wall-traction relation, reaching the reported value would require the assembly to behave as a rotlet sphere of radius 122–277 nm in the 359.2 nm cavity, which at 0.2 Hz is 77% of the cavity radius itself. A rod of half-length 125 nm and half-width 12.5 nm cannot do that. Because the kinematics and the cavity scaling both agree, the disagreement sits in the stress-evaluation step, not in the flow field. As a hypothesis about the origin of the published magnitude rather than a finding, ignoring the 1/r^3^ decay of the rotlet and estimating a linear gradient across the gap gives 0.95 Pa at 1 Hz; that estimate does not reproduce the published frequency dependence, and no mesh-convergence data are available to test it.

Our evaluation is semi-analytic Stokes flow: the Chwang and Wu ellipsoidal friction torque of the source’s own supplementary equations, fed into the exact rotlet-in-a-cavity wall traction, cross-checked against a bead-chain Rotne–Prager–Yamakawa calculation. It **is not a reproduction of a finite-element solution**; the source used a commercial finite-volume solver and we have no equivalent; an independent finite-element treatment of the same rotating-wire problem has since been published for the cytoplasmic case [36]. The published assembly-size sequence also depends on an object detector with a stated precision of 52.2, and the finite-element radii are not the measured means (C3).

#### Level 4, Cox 2016, PIEZO1 gating tension: passes with no tuning, and corrects three things [13]

Seven of eight criteria pass; the single failure is the source’s own internal contradiction (C4). **4.5 and 5.1 mN m**^−1^ **are both cell-attached values**, obtained by two routes in the same paper: 0.4 ×11.3 = 4.52 from the MscL co-expression ratio, and 5.1 from imaging geometry. They are not the two ends of a cell-attached-to-bleb interval; that paper never reports a bleb half-activation tension. The long-standing discrepancy with the 1.4–2.7 mN m^−1^ of Lewis and Grandl [14] is **not** attributable to the pressure-to-tension conversion or to curvature sign: both laboratories use T = Pr/2 and image r directly, the back-inferred dome radii differ by only 1.39-fold, and half-activation pressures on the convex and concave sides are comparable, 16.7 against +19.3 mmHg. The difference lives in the applied pressure, 44 against −16.7 mmHg, and therefore in resting tension and cortical load-sharing. That is why the cell-attached 5.1±0.2 agrees with the *inside-out* 4.7±0.3 (|z| = 1.11) rather than with the cell-attached 2.7 (|z| = 10.7). Neither laboratory is wrong. Because the Laplace relation yields an upper bound, as those authors state twice, **5.1 mN m**^−1^ **is a ceiling, not a central value**.

### 3.5 The dose plane, computed twice

The falsifiable prediction of the placement hypothesis is a dose statement. Using the hemolysis power law calibrated on the plasma-membrane anchor condition, D_plasma = 4.640 × 10^4^, the iso-dose contour through that anchor requires 9.368 Pa at 900 s (figure 7*a*). Because level 3 disagrees with its source, the comparison is computed twice (table 1; figure 7*b*).

**Table 1.** The dose comparison under each candidate value of the intralysosomal wall shear stress.

|  | with the published 1.03 Pa | with our 0.0201 Pa |
|---|---|---|
| Dose ratio D_plasma / D_lysosome | **207**× | **2.80 × 10<sup>6</sup>**× |
| Fragility factor, same-duration caliber | **9.09**× | **466**× |

**Figure 7.**
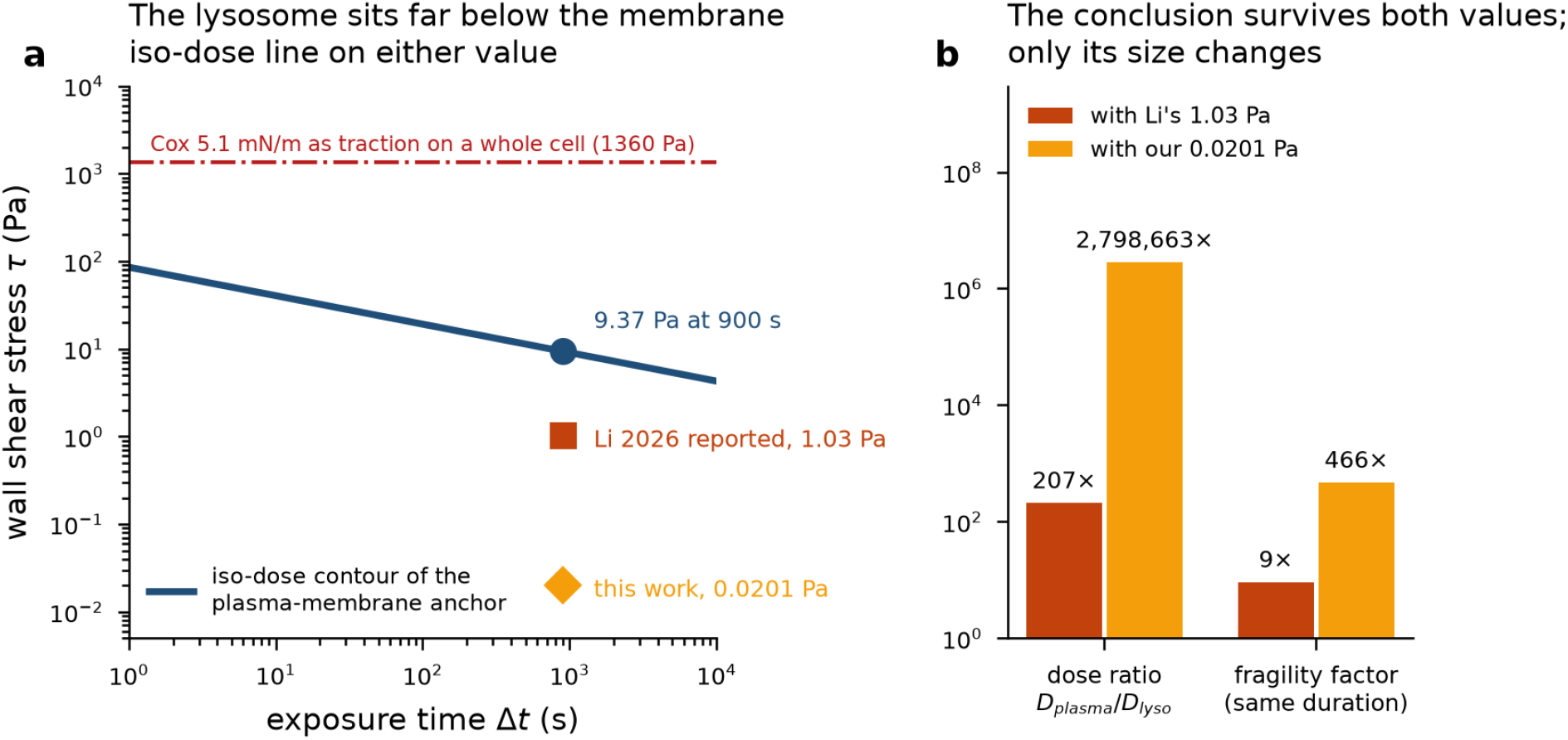
The dose plane, computed twice. **(a)** Iso-dose contour of the plasma-membrane anchor on the stress–time plane, with the two candidate lysosomal wall-shear values at 900 s and the Cox threshold converted to a whole-cell traction. **(b)** Dose ratio and same-duration fragility factor under each value. The conclusion holds under both; only its size changes.

The conclusion is the same under both and only its size changes: for the intralysosomal mechanism to work as described, **the lysosomal limiting membrane must be the weaker membrane**, by between 9 and 466 fold. Since the rupture tension of that membrane has never been measured, no published datum discharges either version. The same-duration fragility factor is the version an experiment can address directly.

Converting the gating threshold into a traction through *τ* = 2T/R gives a result independent of which shear value is adopted: 2.84 × 10^4^ Pa on a lysosome and 1360 Pa on a whole cell. Both are three to five orders of magnitude above any fluid shear either route generates. **Fluid shear cannot open a tension-gated channel at these scales; only anchored traction can.** That inequality predicts an unanchored extracellular swimmer to be mechanically inert at the plasma membrane, which is the regime of the one published experiment that moves cells in the opposite direction.

### 3.6 Response direction as a model output, and what the published record can test

Table 2 sets out the hypothesis operationally, case by case. Four features carry the argument.

**Table 2.**
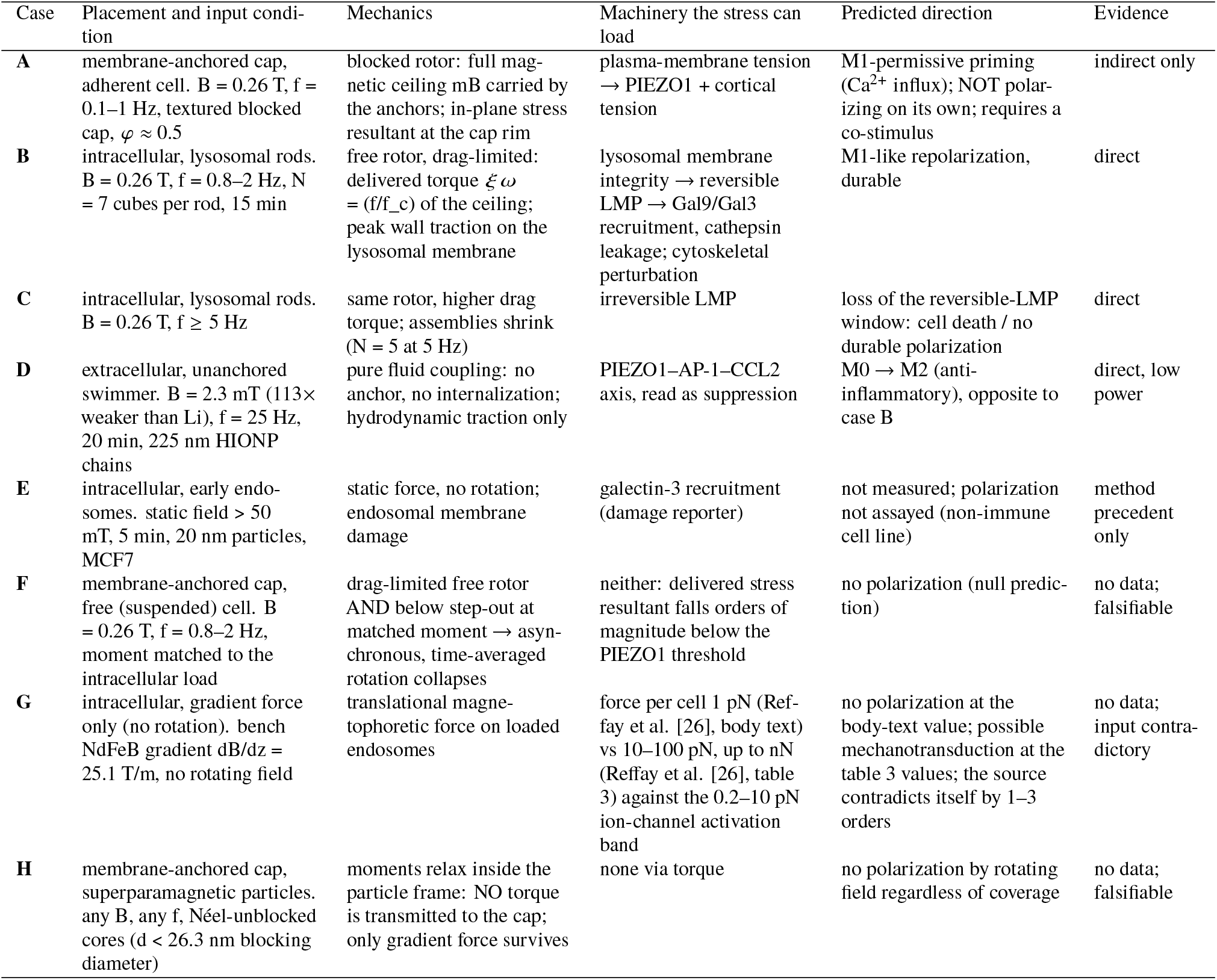
Polarization direction as a model output. Cases B and D are the two published experiments that point in opposite directions; cases F and H are zero-predictions that a single experiment would falsify. The evidence behind each class, with sources, is in electronic supplementary material, table S6.

#### The two contradictory experiments fall out in different rows

Case B, intralysosomal rods at 0.26 T and 0.8–2 Hz, is a free drag-limited rotor whose peak wall traction loads the lysosomal limiting membrane, and durable M1-like repolarization is what is reported. Case D, an unanchored extracellular swimmer at roughly 2.3 mT, delivers a stress resultant of 2.6 ×10^−7^ mN m^−1^ at a step-out margin of 5.4 ×10^−5^, so it is mechanically negligible at the membrane and desynchronized by four decades. Whatever it does, it does not do it by gating PIEZO1, consistent with a reported M2 shift accompanied by *suppression* of that axis.

#### Case A is the honest weak point

A cap on an adherent cell is torque-limited and loads the plasma membrane, but the only evidence is indirect [37], and no magnetic membrane-cap transcriptomic experiment exists.

#### Cases F and H are zero-predictions

(figure 8*b*). A cap on a *free, suspended* cell at matched moment is drag-limited and below step-out, so delivered stress falls orders of magnitude below threshold; a superparamagnetic cap transmits no torque at any coverage, whatever the dose. Each is falsified by one experiment and neither has been attempted.

**Figure 8.**
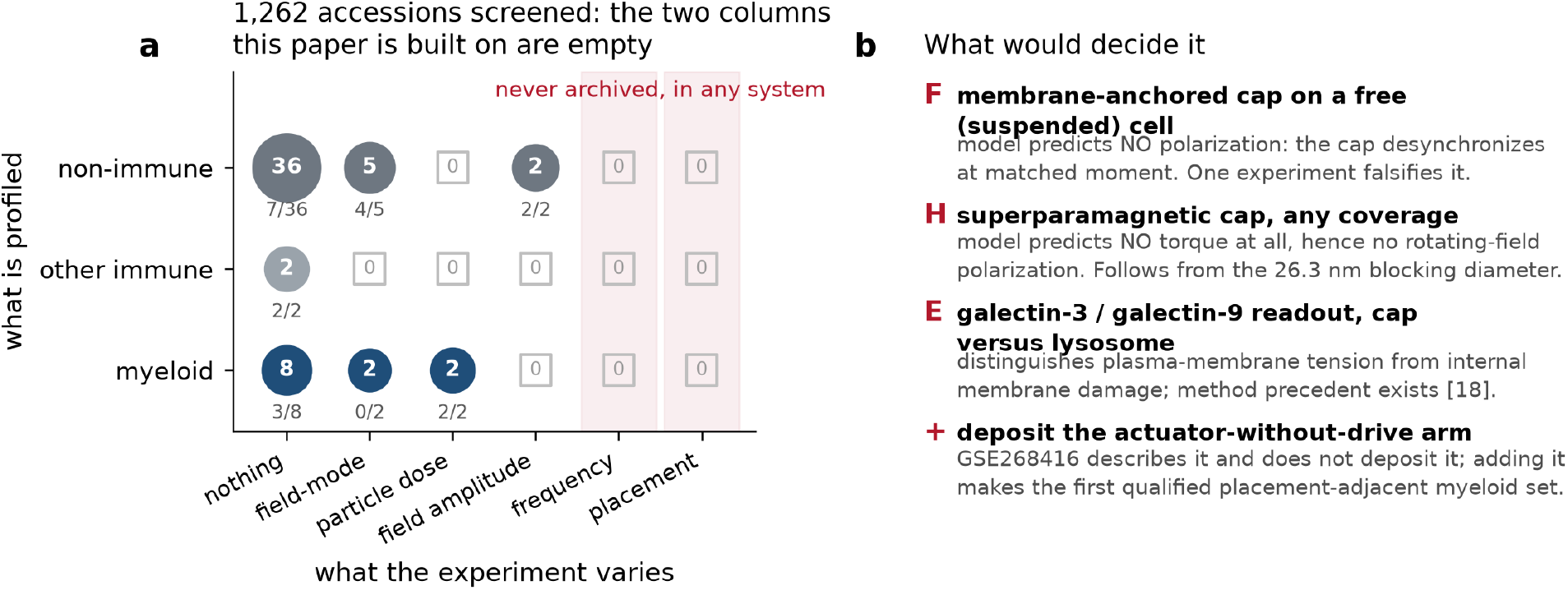
The axis this paper is built on has never been archived. **(a)** Archive map of 1,262 screened accessions: what the experiment varies against what is profiled. Circle area scales with the median n per arm and with the number of datasets; the fraction carrying a processed matrix is printed below each circle; open squares are empty cells. The frequency and placement columns are empty in every row. Circle sizes should be read with the caveat that 34 of 57 mapped rows carry an inferred or unstated n per arm. **(b)** The experiments that would decide it, keyed to the cases of table 2.

Whether archived transcriptomes can adjudicate any of this is itself answerable. Screening the Gene Expression Omnibus (GEO) and ArrayExpress with inclusion rules frozen before the first query returned 1258 unique GEO series and 458 ArrayExpress accessions across 31 queries, four of the latter having no GEO counterpart, for a screened inventory of 1262; 236 series passed a deterministic keyword pre-screen and were read individually (electronic supplementary material, table S5). **No dataset in either archive varies particle placement**, in any cell type, and no myeloid dataset varies drive frequency or field amplitude (figure 8*a*).

Three non-immune datasets do have the designs the field lacks, and they show what such designs recover (figure 9). In the only particle-by-field factorial in the screen [38], the particle term alone moves **63** genes, the field term alone **218**, and the **interaction 1102**, which is 17.5 times the particle term and 5.1 times the field term (figure 9*a*); every positive set-level effect sits in the interaction (figure 9*b*). A three-arm design cannot estimate that interaction and a two-arm design estimates neither term separately, which is what nine of the twelve core myeloid datasets are. In a second dataset carrying particles in *both* field arms [39,40], a 50 T m^−1^ gradient produces **no detectable transcriptional change in the cell body** and **1278 changes in the neurite of the same cells** (figure 9*c*): a force present everywhere can produce a readout confined to one compartment, so a whole-cell measurement scores a genuine mechanical input as a null. In a third [41], tripling a magnetically delivered strain from 5% to 15% buys only 1.22 times the score and 1.18 times the gene count (figure 9*d*), so a single high-amplitude arm is a poor probe of the dose axis.

**Figure 9.**
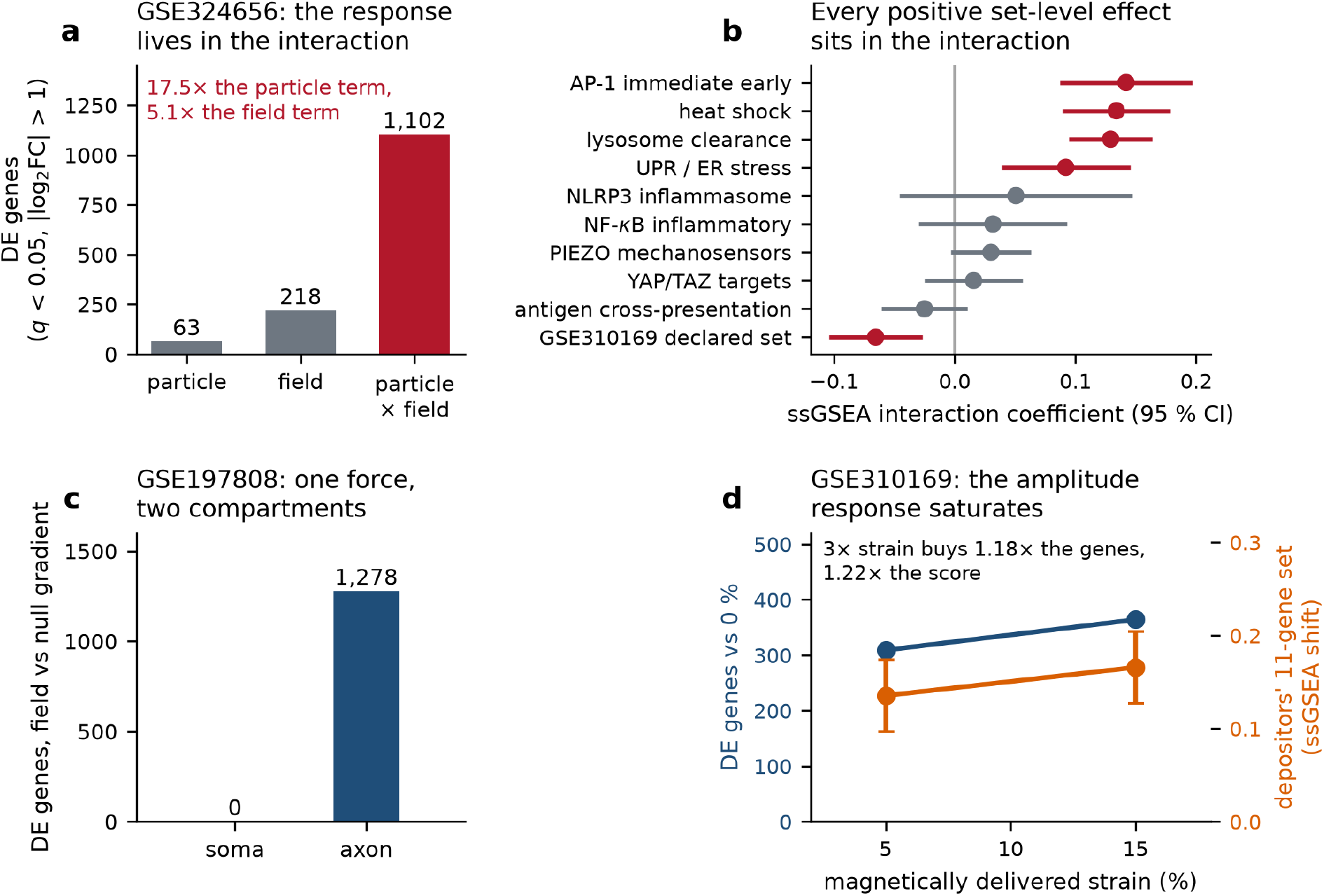
What a proper design recovers, in systems that have run one. **(a)** GSE324656, the only particle ×field factorial in the screen: differentially expressed genes attributable to each term. **(b)** Set-level interaction coefficients with 95 % confidence intervals; red, q < 0.05. Every positive effect is in the interaction. **(c)** GSE197808: the same field applied to particle-loaded neurons, read in two compartments. **(d)** GSE310169: gene count (left axis) and the depositors’ own eleven-gene set (right axis, 95 % CI) across graded magnetically delivered strain.

## 4 Discussion

Two placements of the same particle, matched on the quantity the field reports, are two different machines. Their step-out frequencies scale with assembly size in opposite directions and cross inside the working window of published experiments, so an experimenter who adds particles improves one actuator and degrades the other. Their torque economies differ by three orders of magnitude, and adhesion switches the membrane route between regimes while leaving the intracellular route untouched. Their tension resultants are governed by different variables: the cap by the magnetic texture of its monolayer, with cell radius canceling exactly, the rod by the viscosity caliber of its surroundings, with cavity radius almost irrelevant. None of this is visible if placement is treated as a synthesis detail.

The practical reading is a short list of design rules. Report magnetic texture, not only particle size and field, because random easy axes cost a factor of 808 and superparamagnetic caps cost everything. State whether cells were adherent or suspended during actuation, because that one word decides whether a membrane cap is torque-limited or drag-limited. Do not scale an intracellular actuator by adding particles. Do not transfer a frequency chosen for one placement to the other, since the crossing at 4.04 Hz sits inside the window both use.

The model’s weakest joints are the ones we have named rather than smoothed. Two inputs have never been measured, and each bounds a central claim: without the lysosomal rupture tension the dose argument reduces to a requirement on an unmeasured number, and without the plasma-membrane screening length the cap prediction reverses sign across the plausible range. The cortical load-sharing fraction is contradictory within one source by a factor of 2.67, and it decides which side of the PIEZO1 threshold a conventional Janus cap falls on. Our level-3 evaluation disagrees with its source by 33 to 81 fold in magnitude while reproducing its structure, and we could not test the hypothesis that the gap lies in the stress-evaluation step, because the source reports no mesh-convergence data and we had no equivalent solver. The membrane arm is analytic throughout; we do not compute stress fields, channel open probabilities, or the correction that cortical coupling would apply to the threshold. The immunological direction is predicted, not measured: no magnetic membrane-cap transcriptomic experiment exists, and the quantitative readouts we report come from non-immune systems chosen because they ran designs the myeloid literature has not.

What would settle it is modest, and figure 8*b* keys each missing experiment to the case it would decide. Case F and case H are zero-predictions, each falsified by a single experiment that no one has run: a matched-moment cap on a suspended cell should produce nothing, and a superparamagnetic cap should produce nothing at any coverage or field. A particle-by-field factorial in macrophages, with the actuator-without-drive arm that nine of twelve core datasets omit, would supply the interaction term that the one factorial in the archive shows to be five to seventeen times larger than either main effect. Measuring the lysosomal rupture tension would convert the dose argument from a requirement into a test. Until then, the proposition stands as a model with its bounds marked: where the magnet sits sets the sign of the scaling law, and the biology follows the mechanics rather than the dose.

## 5 Methods

All quantities are computed in closed form or by numerical integration of ordinary differential equations; no parameter is fitted to any dataset the model is tested against, and no experiment was performed. Scripts, inputs and outputs are available as described below; the electronic supplementary material gives the derivations, the frozen pass criteria, and the full provenance of every constant.

Single-particle moments follow from bulk saturation magnetization and particle volume, with the Néel blocking diameter obtained from the standard relaxation criterion at 300 K and a 100 s measurement time. Rod rotational friction is evaluated on two independent calibers, the Broersma slender-cylinder expansion and the Chwang and Wu prolate-spheroid torque, reported side by side; whole-cell drag uses the Stokes sphere. Wall traction inside a spherical cavity uses the exact axisymmetric rotlet solution with the finite-size correction of the source, cross-checked against a bead-chain Rotne–Prager–Yamakawa mobility calculation [42,43]. Rotation dynamics are obtained by integrating the lag-angle equation for the permanent-moment and induced-moment branches separately with an adaptive solver, and verified against the closed-form mean rotation rate. Cap mechanics follow from reacting the magnetic torque as an in-plane stress resultant at the cap rim, with tension propagation computed cortex-free and screened. Dose comparisons use a hemolysis-calibrated power law anchored on the plasma-membrane condition.

Public transcriptome screening used inclusion rules written and frozen before the first query. Differential expression used PyDESeq2 [44] with main and interaction coefficients estimated in a single factorial model; set-level scores used single-sample gene-set enrichment [45] on log_2_(counts per million + 1) followed by ordinary least squares and Benjamini–Hochberg control. Datasets with fewer than three samples per arm are reported descriptively and carry no statistical claim.

## 6 Data accessibility

All scripts, computed tables and figure source data are available at https://github.com/congzhaoqing/magnetic-particle-placement, release v1.0.4, which is the state accompanying this manuscript. The electronic supplementary material comprises table S1, the provenance of all 142 physical inputs (`Table1_parameter_provenance_merged.csv`); table S2, the pre-registered criteria and measured quantities of all four validation levels (`validation_ladder_summary.csv`); table S3, the five registered source contradictions (`F_source_contradictions.csv`); table S4, the step-out scaling curves and the thirteen matched-moment configurations (`F_two_path_stepout_scaling.csv, F_two_path_matched_moment.csv`); table S5, the screened archive inventory with tier assignments (`F_GEO_scan_inventory.csv`); and table S6, the per-case published evidence behind table 2 (`F_table_S6_case_evidence.csv`). No new experimental data were generated. The transcriptome datasets re-analyzed are public and are identified by accession in the electronic supplementary material.

## 7 Declaration of AI use

Generative artificial intelligence was used to assist with literature screening, code drafting and language editing. All quantitative results were produced by the scripts in that repository, and all numerical claims were checked against those scripts by the authors, who take full responsibility for the content.

## 8 Authors’ contributions

Y.X.: data curation, formal analysis, software, visualization. L.Y.: methodology. A.L.: software, validation. J.T.: visualization. Z.C.: conceptualization, data curation, funding acquisition, methodology, supervision, writing – original draft, writing – review and editing. All authors read and approved the manuscript.

